# Neural correlates of temporal presentness in the precuneus: a crosslinguistic fMRI study based on speech stimuli

**DOI:** 10.1101/2020.06.17.158485

**Authors:** Long Tang, Toshimitsu Takahashi, Tamami Shimada, Masayuki Komachi, Noriko Imanishi, Yuji Nishiyama, Takashi Iida, Yukio Otsu, Shigeru Kitazawa

**Affiliations:** Dynamic Brain Network Laboratory, Graduate School of Frontier Biosciences, Osaka University, Osaka 565-0871, Japan; Department of Brain Physiology, Graduate School of Medicine, Osaka University, Osaka 565-0871, Japan; Center for Information and Neural Networks (CiNet), National Institute of Information and Communications Technology, Osaka 565-0871, Japan; Department of Physiology, Dokkyo Medical University School of Medicine, Tochigi 880, Japan; Faculty of Languages and Cultures, Meikai University, Chiba 279-8550, Japan; Faculty of Humanities and Social Sciences, Shizuoka University, Shizuoka 422-8529, Japan; The University of Tokyo (emeritus), Tokyo 113-0033, Japan; Professor Emeritus, Keio University, Tokyo 108-8345, Japan; Faculty of Foreign Language Studies, Kansai University, Osaka 564-8680, Japan

**Keywords:** cross-linguistic, fMRI, precuneus, tense, time perception

## Abstract

The position of any event in time could be either present, past, or future. This temporal discrimination is vitally important in our daily conversations, but it remains elusive how the human brain distinguishes among the past, present, and future. To address this issue, we searched for neural correlates of presentness, pastness, and futurity, each of which is automatically evoked when we hear sentences such as ‘it is raining now’, ‘it rained yesterday’, or ‘it will rain tomorrow’. Here, we show that sentences that evoked ‘presentness’ activated the bilateral precuneus more strongly than those that evoked ‘pastness’ or ‘futurity’. Interestingly, this contrast was shared across native speakers of Japanese, English, and Chinese, languages which vary considerably in their verb tense systems. The results suggest that the precuneus serves as a key region that provides the origin, the Now, to our time perception irrespective of differences in tense systems across languages.

## Introduction

Each position in time is either in the past, present, or future. This way of discrimination in time, which McTaggart (McTaggart JE 1908) named the *A* series, is essential in our daily conversation. When we hear someone saying “it looks like rain today”, for example, we are instantly aware that the speaker is addressing the weather in the near future. If we say “it is raining in Osaka”, the temporal position of the remark is present. It is evident that the remark “it rained on and off all day” refers to the weather some time in the past. Without reference to time in each remark, we are not able to exchange comments on the weather. It is remarkable that the process of allocating each rainy event to present, past, or future is so automatic that we cannot suppress a feeling of “futurity” arising from the remark “it looks like rain”. Which part of the brain discriminates among “futurity”, “presentness”, and “pastness” without any conscious effort?

It is generally accepted that when we recall some events in the past, portions of the memory system of the brain, such as the hippocampal and parahippocampal network and the inferior frontal gyrus, are recruited (McClelland JL et al. 1995; Okuda J et al. 2003; Addis DR et al. 2007; Szpunar KK et al. 2007; Corkin S 2013; Wilson AG et al. 2013). Interestingly, similar regions are recruited when we consciously imagine some events in the future (Maguire EA et al. 1998; Hassabis D and EA Maguire 2007; Addis DR et al. 2011; Martin VC et al. 2011). These studies suggest that the memory system supports our feelings of pastness as well as futurity.

In a recent functional imaging study, Peer M et al. (2015) reported that the precuneus, inferior parietal cortex, and medial frontal cortex are involved in mental orientation in space, time, and person. These regions overlap with the default mode network, one of a few basic cortical networks that are active even in a resting state. Peer M *et al*. (2015) further reported that some specific parts of the default mode network (anterior part of the precuneus and the posterior part of the medial frontal cortex) are involved in mental orientation in time.

These previous studies seem to highlight the importance of a number of medial cortical systems in representing the *A* series of time. However, previous studies have adopted mental tasks that were carried out with effort. Participants were required to recall actual events in the past or imagine possible events in the future. Furthermore, these tasks did not directly address the issue of presentness that serves as the origin of the *A* series. In the present study, we aimed to elucidate the neural bases of presentness in particular in addition to those of pastness, and futurity that are automatically evoked whenever we hear the speech of others in our daily conversations. For this purpose, we prepared short sentences such as “I eat curry today” or “I ate curry yesterday”, each of which contained a verb and an adverb of time, and examined brain regions whose activity correlated with presentness, pastness, and futurity that were evoked in each participant without any effort.

Developmental linguistic studies have shown that the ability to discriminate between the past, present, and future develops over several years from 2.5 to 6.5 years of age (Weist RM et al. 1991; Busby J and T Suddendorf 2005; Busby-Grant J and T Suddendorf 2009; Tillman KA et al. 2017; Zhang M and JA Hudson 2018, 2018), but the speed of development varies across different languages (Weist RM *et al*. 1991). Therefore, it is crucial to ask whether the neural correlates of the *A* series are shared across speakers of different languages. To maximize the variability across languages, we chose English and Chinese in addition to Japanese. The tense of the verb, which is taken for granted in European languages such as English, could be absent in many non-European languages such as Chinese. To find neural bases that are shared across speakers of different language systems, we carried out experiments with speakers of these three languages. Here, we show that the precuneus, a core region of the default mode network, serves as a cross-linguistic core region that responded two to three times greater to presentness than to pastness or futurity.

## Materials and methods

### Participants

Fifty-four healthy young adults (32 males and 22 females ranging in age from 20 to 38 years) participated in the experiment. The participants were either native speakers of Japanese (n=18), English (n=18), or Chinese (n=18). All participants had normal hearing, with no history of neurological disorders and were naïve to the purpose of the experiments. They were all strongly right-handed according to the Edinburgh Inventory (Oldfield RC 1971). This study was approved by the Ethical Review Board of Osaka University, Graduate School of Frontier Biosciences. Written informed consent was obtained from all participants before the experiments.

### Stimuli

We initially prepared 180 sentences in Japanese. Each sentence consisted of three parts: an adverb of time, a noun with a particle, and a verb. Let us take the sentence “昨日、本を読んだ ∼ I read books yesterday”, for example. It started with an adverb that pointed to the past (昨日= yesterday), which was followed by a noun with a particle (本を = book) and a final verb in the past tense (読んだ = read [réd]).

Of the 180 sentences, 144 were grammatically correct. We chose from ten adverbs of time: three related to the past (先月、先週、昨日; last month, last week, yesterday), four to the present (今、今日、今週、今月; now, today, this week, this month), and three to the future (明日、来週、来月; tomorrow, next week, next month). We chose one noun + verb pair from 30. Each verb was either in the present form (e.g., 読む = read [ri:d]) or in the past tense with the help of the auxiliary verb (読んだ= read [réd]). We thus prepared 600 grammatically correct sentences in total (10 adverbs × 30 noun+verb × 2 tenses). We chose 144 from the 600. We also prepared 36 “nonsensical” sentences. They were illogical either in its combination of the noun and the verb (e.g., I ate the road today) or in its combination of the adverb and the tense of the verb (e.g., I washed my car next week). These stimuli were grouped into six sets (n = 30), and each set consisted of 24 correct and 6 incorrect sentences (Table 1).

**Table 1.** Examples of grammatical and nonsensical sentences in three languages.

| No. | Japanese | English | Chinese |
| --- | --- | --- | --- |
| 1 | 今日手紙を送る | I send a letter today | 今天我寄信 |
| 2 | 今週実家を出た | I left my parents' house this week | 这周我离开了老家 |
| 3* | 昨日進学を決める | I decide on continuing school<br>yesterday | 昨天我决定升学 |
| 4 | 来週バスに乗る | I will ride a bus next week | 下周我坐公交车 |
| 5* | 今日天井を学ぶ | I will study ceilings today. | 今天我学天花板 |
| 6 | 先月セールを開始した | I started a sale last month | 上个月我开始了大甩卖 |
| 7 | 先月カギを紛失した | I lost my key last month | 上个月我丢失了钥匙 |
| 8 | 来月開催を決定する | I decide on starting the fair next<br>month | 下个月我决定召开会议 |
| 9 | 今週財布を落とす | I will drop my wallet this week | 这周我掉钱包 |
| 10 | 今月新メニューを作った | I made a new menu this month | 这个月我制作了新菜单 |
| 11 | 来週宝物を発見する | I will discover treasure next week | 下周我发现财宝 |
| 12 | 来週財布を失くす | I will lose my wallet next week | 下周我丢失钱包 |
| 13 | 今会議を始める | I will start the meeting | 现在我要开始会议 |
| 14 | 今月ペンキを塗る | I paint it this month | 这个月我涂油漆 |
| 15 | 明日避難場所を見つける | I will find the evacuation shelter<br>tomorrow | 明天我要发现避难场所 |
| 16 | 今週東京に着く | I arrive in Tokyo this week | 这周我到达东京 |
| 17 | 今ガスを止めた | I stopped the gas now | 现在我关闭了煤气 |
| 18 | 昨日日記を書いた | I wrote my journal yesterday | 昨天我写了日记 |
| 19 | 先月会場に集合した | We gathered at the site last month | 上个月我们聚集在了会场 |
| 20* | 昨日踊りを消した | I erased the dance yesterday | 昨天我关了舞蹈 |
| 21* | 先週卒業に集まる | We gather at the graduation last week | 上周我们聚集在毕业 |
| 22* | 来週愛車を洗った | I washed my car next week | 下周我洗了车 |
| 23 | 今日カレーを食べた | I ate curry today | 今天我吃了咖喱饭 |
| 24 | 今月テントを乾かす | I will dry the tent this month | 这个月我要晒干帐篷 |
| 25 | 今週ポスターを貼った | I posted the poster this week | 这周我贴了海报 |
| 26* | 今月発表をあげた | I raised a presentation this month | 这个月我给了你发表 |
| 27 | 今月日本を出発した | I left Japan this month | 这个月我离开了日本 |
| 28 | 先月アフリカに到着した | I arrived in Africa last month | 上个月我到达了非洲 |
| 29 | 明日本を読む | I will read books tomorrow | 明天我读书 |
| 30 | 先週商品を売った | I sold the merchandise last week | 上周我卖了商品 |

Asterisks show nonsensical sentences. Thirty sentences in one of the six sets of sentences are listed.

The 180 Japanese sentences were then translated into English and Chinese sentences by native speakers. Linguists of each language edited the translations to make the grammatical sentences as natural as possible while maintaining the original pastness, presentness, and futurity that each original Japanese sentence evoked. For example, the adverb of time was placed at the top in the Chinese translation but moved to the end in English translation. However, to make the duration of each sentence as constant as possible, we avoided using present progressive forms in English.

Each sentence was read aloud by female native speakers of each language. The female voices were sampled at 44100 Hz and recorded digitally in a “wav” format. The duration of each stimulus was approximately 2 s (2.2 ± 0.4 s, mean ± s.d.).

### Task procedures in the scanner

The participants lay supine in a 3T magnetic resonance (MR) scanner (MAGNETOM Trio, Siemens, Munich, Germany) with earphones in the ears and with their eyes closed. During each scanning session (360 s), thirty sentence stimuli were presented one by one with an intertrial interval of 12 s (Fig. 2A). The participants were asked to judge whether they felt each sentence natural or unnatural. When they felt that a sentence was natural, they did nothing. Otherwise, they responded by pressing a button after a beep sound (0.5 s) that was always presented at 6 s after the onset of each narration. They completed six sessions with six sets of sentences and made a natural/unnatural judgment to each of the 180 sentences. The presentation of the auditory stimuli and detection of button presses were controlled by Presentation Software (Neurobehavioral System, San Francisco, CA).

**Figure 1.**
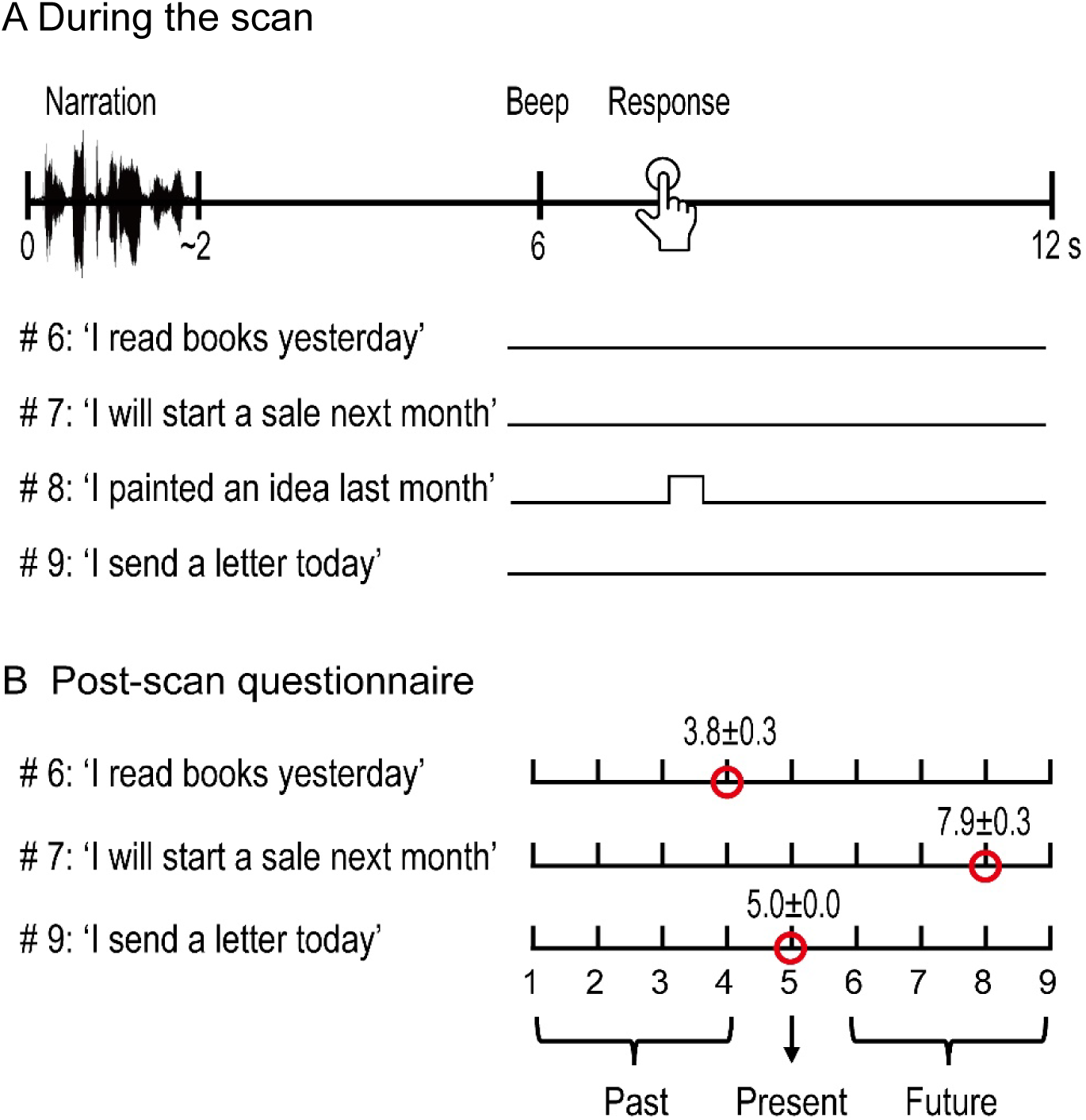
Experimental procedures during (A) and after (B) the scan. (A) During the MRI scan, a short narration (∼2 s) was presented, which was followed by a beep at 6 s. After the beep, the participants pressed a button when they felt the sentence was unnatural. A typical participant made no response to stimuli #6, #7, and #9 but responded to the unnatural sentence #8 (I painted an idea last month). (B) After the scan, the participant rated each sentence that was presented in the scanner on a nine-point scale (1: the far past…, 5: present…, 9: far future). The sentences without responses were classified into ‘past’ (1 to 4), ‘present’ (5), and ‘future’ (6 to 9) groups. The participants rated #6, 7, and 8 as near past (4), future (8), and present (5). The numbers show the mean rating across participants (n = 18) with the standard error of the mean.

**Figure 2.**
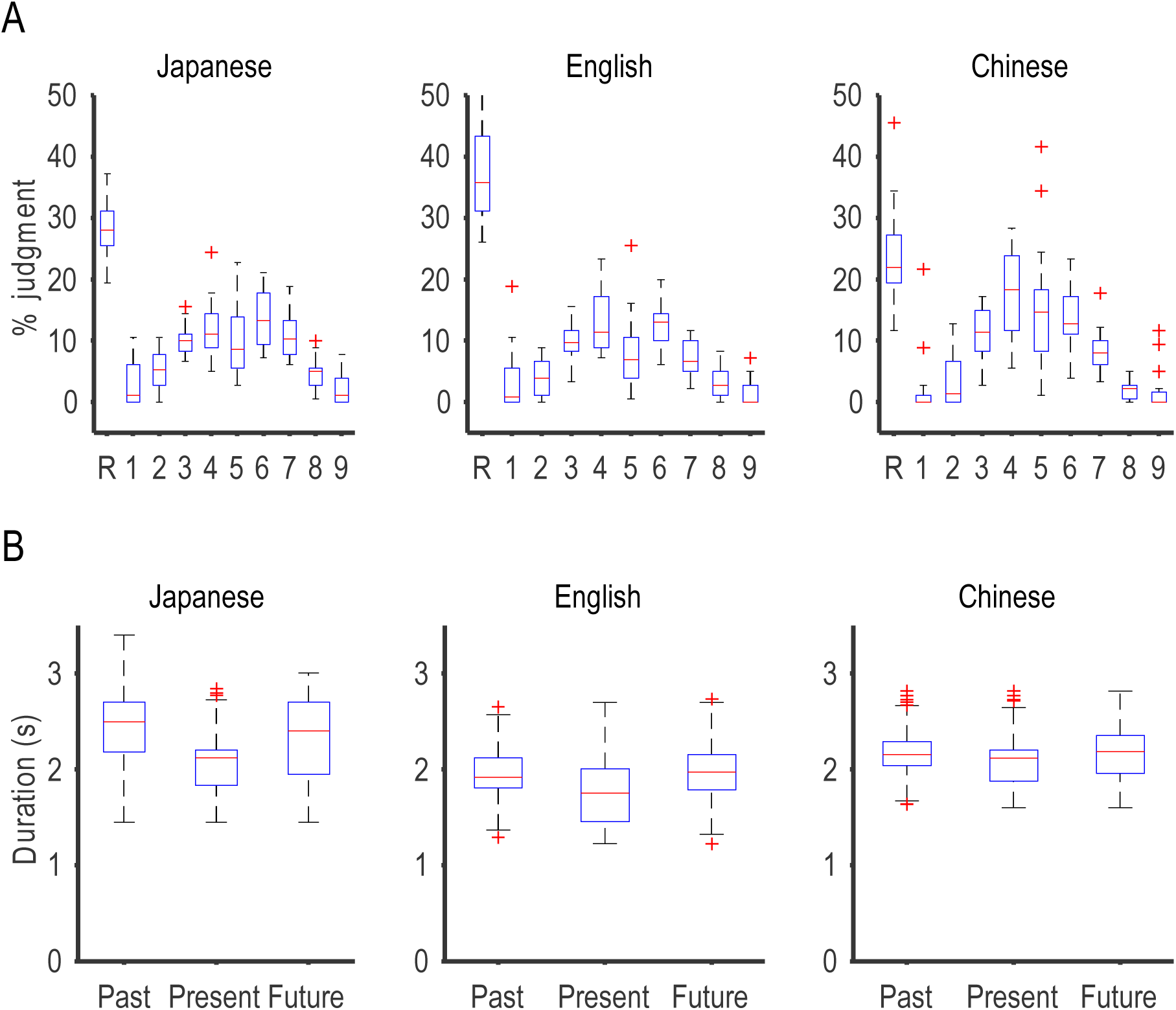
Behavioral results. (A) Percentage of sentences that were judged as unnatural during scan (response; R) and rated on a nine-point scale (1-9) after the scan. Each panel shows data from participants of the same language (n = 18 for each panel). Box plots show the median (red bar) and 25th and 75th percentiles (box). Outliers are shown by +.(B) Durations of sentences that were judged as natural in each language. Note that the mean duration of the present sentences was significantly shorter than the others (p<.001, post hoc t-tests). Note also that the mean duration of Japanese sentences was significantly longer than the others.

Functional images sensitive to the blood oxygen level-dependent (BOLD) contrast (Ogawa et al. 1993) were obtained from a gradient-echo echo-planar imaging pulse sequence with a 192-mm field-of-view, 4-mm slice thickness, 1-mm interslice gap, a 64 × 64 data matrix, a repetition time (TR) of 2000 ms, an echo time (TE) of 30 ms, and a flip angle of 70°. The image volumes covered the entire brain with 30 slices. High-resolution T1-weighted anatomical images were also acquired for each participant using the 3-D MP-RAGE pulse sequence with TR of 2250 ms, TE of 53 ms, flip angle of 59°, and voxel size of 1 × 1 × 1 mm3.

### Postscan evaluation of time

It was not until after MRI measurements were completed that the participants evaluated the time that each sentence referred to. They rated their temporal evaluation by using a nine-point scale: they marked 5 when a sentence felt like it had referred to the “present”, marked 1 when it felt “far past”, and marked 9 when it felt “far future” (Fig. 2B). We defined three groups according to these ratings: past (1-4), present (5), and future (6-9) groups (Fig. 2B).

### Data Analysis

Brain activations of 54 participants (18 native speakers of Japanese, 18 native speakers of Chinese, and 18 native speakers of English) were analyzed by using SPM12 (Wellcome Department of Cognitive Neurology, London, UK) implemented with MATLAB R2018a (MathWorks, Natick, MA, USA). The two main effects of time (past, present, and future; within-subject) and language (Japanese, English, and Chinese; between-subject) and their interaction were the major targets of data analysis. We also analyzed the effect of naturality (with/without button press).

The data preprocessing for each experimental session proceeded in the following 5 steps: motion correction, slice timing adjustment, coregistration of the anatomical T1 images with the mean functional images in a session, spatial normalization of all images to the Montreal Neurological Institute (MNI) reference brain in 2-mm isovoxel resolution, and spatial smoothing with a Gaussian kernel of 8-mm full-width at half-maximum. The data were further analyzed by using a conventional 2-stage random effect model (Büchel C et al. 1998).

In the first-level analysis, six sessions of data from each participant were combined, and a linear model with the following 26 regressors (6 x 4 finite impulse response, FIR, bases and one hemodynamic function) was fitted to the data from each participant. Six FIR bases with an impulse width of 2 s were prepared for each of the past trials at 0, 2, 4, 6, 8, and 10 s from each stimulation onset. Similarly, 6 FIR bases were prepared for each of the present, future, and response trials. To regress out the effect of the beep sound and the button response, two more regressors with a conventional hemodynamic function were added to the model. To obtain a set of empirical hemodynamic responses at the six time points, we aligned MRI signals at speech onset, averaged the signals over all gray matter voxels for each participant, normalized them by the standard deviation (z-score transformation), and finally averaged across the 54 participants. We thus obtained a set of empirical hemodynamic responses of [-0.055, -0.11, 0.083, 0.13, -0.027, -0.040] at 0, 2, 4, 6, 8, and 10 s. The empirical response curve showed an initial dip at 2 s, peaked at 6 s, and returned quickly to the baseline toward 10 s. To obtain a beta value for the past trials, for example, we took an inner product between the response vector and each coefficient vector that consisted of six coefficients estimated for each of the six FIR bases prepared for the past trials. We thus obtained four beta values, one for each of the past, present, future, and response trials. These beta values were transferred to the second-level analyses.

In the second-level analyses, we carried out the following four analyses and obtained four statistical maps: (1) one-way ANOVA (within-subjects) for the main effect of time (past, present, and future), (2) one-way ANOVA (between-subjects) for the main effect of language (Japanese, English, Chinese), (3) two-way mixed model ANOVA for the interaction between time and language, and (4) one-way ANOVA (within-subjects) for the main effect of response. To examine the effect of stimulus duration, we conducted one-way ANOVA (within-subjects) for the main effect of stimulus duration (short, middle, and long). Each statistical map was initially thresholded at a P-value of 0.005 (voxel level, uncorrected). For each cluster of voxels that satisfied this criterion, the chance of finding a cluster with this or a greater size within the search volume was calculated once uncorrected (cluster-level P, uncorrected) and further corrected for the familywise error rate [cluster-level P, familywise error rate (FWE)-corrected] as implemented in SPM12. We set a threshold of 0.05 and identified clusters that satisfied the FWE-corrected criterion (cluster P < 0.05, FWE-corrected). Furthermore, we identified clusters that contained any voxels that met a peak level FWE-corrected criterion (peak level P < 0.05, FWE-corrected). After one-way ANOVA (within-subjects) for the main effect of time, we repeated one-sample t-tests as post hoc tests (past < present, past < future, present < future, past > present, past > future, and present > future).

## Results

### Behaviors

In a 3 Tesla MRI scanner, each participant listened to a series of short narrations (∼2 s) that was presented one by one with an intertrial interval of 12 s (Fig. 1). Each narration in English, for example, started with “I”, followed by a verb (e.g., read [réd]), an object (books), and an adverb of time (yesterday). The participants were required to press a button only when they felt a sentence was unnatural (e.g., I painted an idea last month). We presented nonsensical sentences in 20% of the entire trials. The fifty-four participants pressed the response button in 30% (n = 2887) of the total trials (n = 9720). In the remaining 70% of the trials, the participants made no response.

Those trials with no response were classified after the scan into past (32%, 3109), present (11%, 1080), and future trials (27%, 2644), according to the postscan rating on a 9-point scale questionnaire (Fig. 2A). It is worth noting that participants were not told to judge pastness, presentness, or futurity in the scanner. We adopted this postscan rating design because we wanted to find neural correlates of the feeling of pastness, presentness, and futurity that were automatically evoked, as occurs our daily conversations.

Two-way analysis of variance with two main effects of the classification (a response or a rating from 1-9) and language (Japanese, English, Chinese) showed that the interaction was highly significant (F(18, 459) = 5.9, p < 0.0001). Post hoc t-tests (simple main effects) showed that 1) the English group yielded a greater percentage of ‘unnatural’ judgments (R) than the other groups (p < 0.0001), and 2) the Chinese group yielded greater percentages of ‘near past’ (rating 4, p < 0.01) and ‘present’ (rating 5, p<0.01) judgments than the other groups.

We planned to make the duration of voice stimuli as identical as possible approximately 2 s. They did distribute near 2 s, but a two-way ANOVA (time x language) revealed that there were some significant differences (Fig. 2B). The main effect of language was highly significant (F(2, 6824) = 699, p < 0.0001). The mean duration descended in the order of Japanese (2.3 s), Chinese (2.1 s), and English (1.9 s). The main effect of time classification was also highly significant (F(2, 6824) = 226, p < 0.0001). The mean duration descended in the order of the past (2.18 s), future (2.15 s), and present (1.98 s).

### fMRI

The main effect of time (past, present, and future) was significant in three clusters after familywise error corrections (Table 2). These clusters were located in the bilateral precuneus (Fig. 3A), in the left Broca’s area (Fig. 3B), and in the left transverse gyrus of Heschl (Fig. 3C). The mean response curves in the precuneus cluster showed that the region responded to the present stimuli 2-3 times stronger than the future or past stimuli. Indeed, post hoc tests showed that the response to the present stimuli dominated over the other two categories of stimuli (Fig. 3G). The second cluster in the left Broca’s area also showed the greatest activations to the present stimuli (Figs. 3E, H). On the other hand, responses in the left transverse gyrus of Heschl were weakest with exposure to the present stimuli (Figs. 3F, I).

**Table 2.**
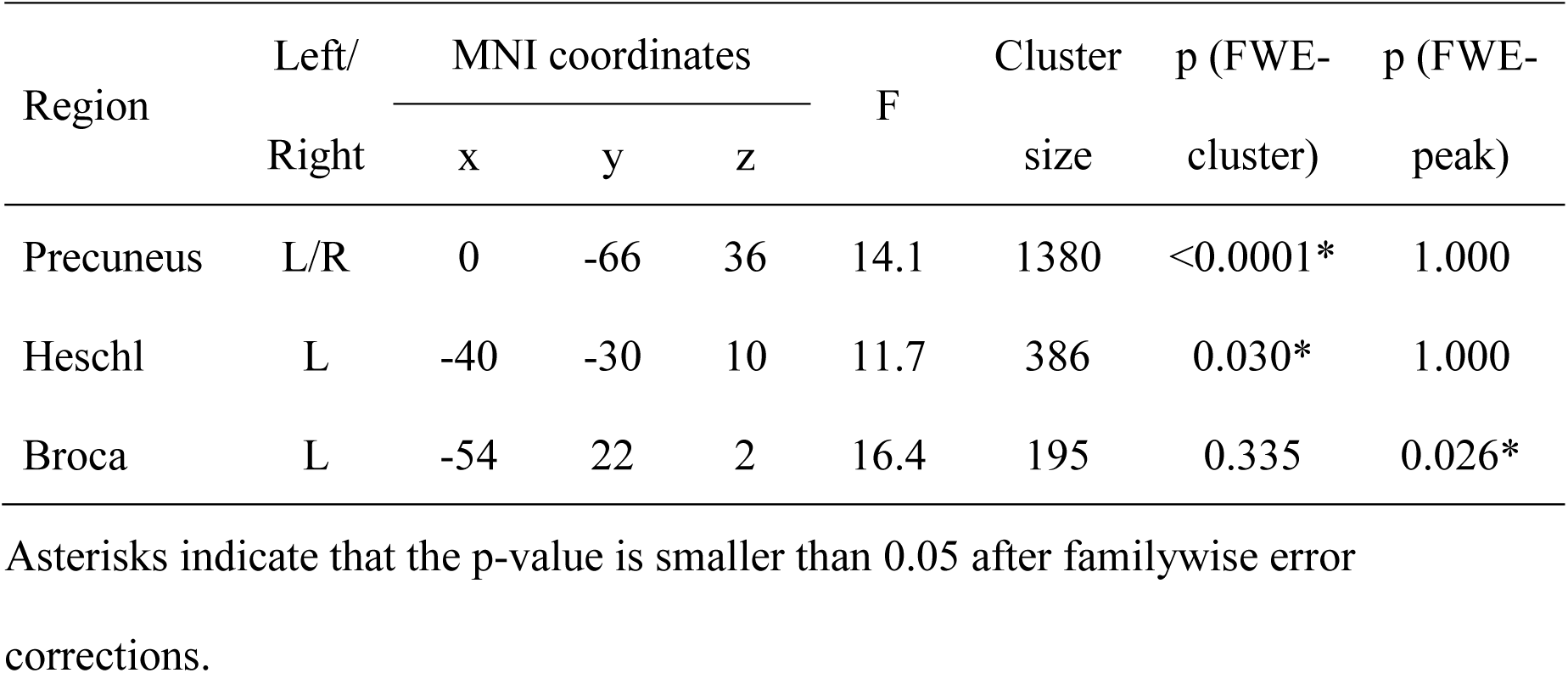
Brain regions that showed a significant main effect of time.

| Region | Left/<br>Right | MNI coordinates |  |  | F | Cluster<br>size | p (FWE-cluster) | p (FWE-peak) |
| --- | --- | --- | --- | --- | --- | --- | --- | --- |
|  |  | x | y | z |  |  |  |  |
| Precuneus | L/R | 0 | -66 | 36 | 14.1 | 1380 | <0.0001* | 1.000 |
| Heschl | L | -40 | -30 | 10 | 11.7 | 386 | 0.030* | 1.000 |
| Broca | L | -54 | 22 | 2 | 16.4 | 195 | 0.335 | 0.026* |

Asterisks indicate that the p-value is smaller than 0.05 after familywise error corrections.

**Figure 3.**
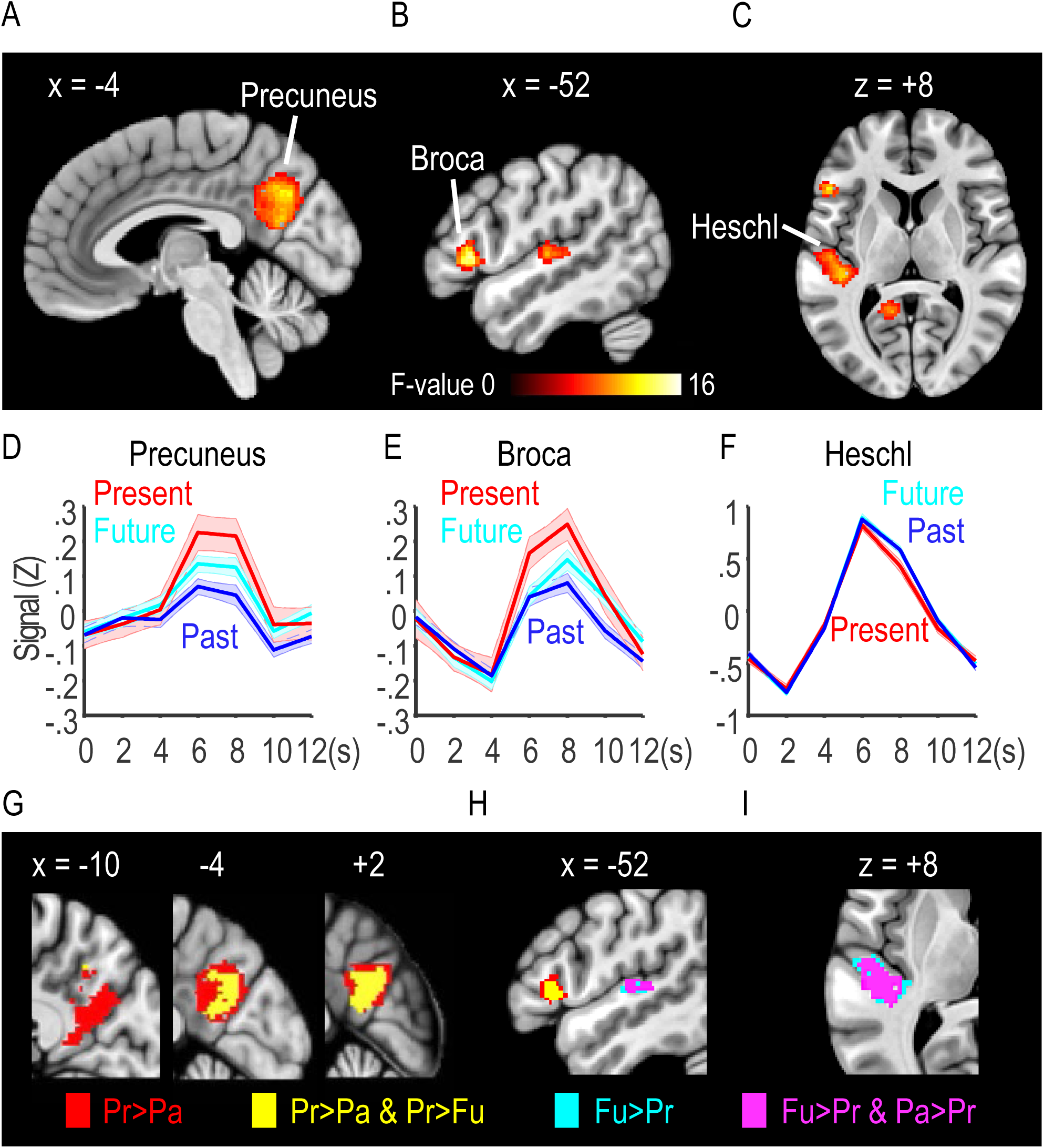
Three clusters that showed a significant effect of time (past, present, and future). (A-C) Clusters in the precuneus (A), Broca’s area (B), and transverse gyrus of Heschl (C) survived after familywise error corrections (p < 0.05). (D-F) The mean responses (z-scores) in the three clusters after the onset of the present (red), future (green), and past (blue) stimuli. Shaded regions show the s.e.m. across 54 participants. (G-I) Results of the six post hoc comparisons. Four of them (Present>Past, Present>Future, Future>Present, Past>Present) yielded significant differences (p < 0.005, uncorrected). Different colors show different combinations shown at the bottom.

The main effect of language (Japanese, English, and Chinese) was significant in one cluster that was centered in the left lingual gyrus (Table 3, Fig. 4A). Post hoc analyses showed that Japanese participants showed much greater responses than the other two groups of participants (Fig. 4B). Analyses of the interaction between the main effects of time and language yielded no regions after the familywise error corrections.

**Table 3.** Brain regions that showed a significant main effect of language.

| Region | Left/<br>Right | MNI coordinates |  |  | F | Cluster<br>size | p (FWE-cluster) | p (FWE-peak) |
| --- | --- | --- | --- | --- | --- | --- | --- | --- |
|  |  | x | y | z |  |  |  |  |
| Lingual<br>gyrus | L/R | -12 | -80 | -2 | 25.0 | 3652 | <0.0001* | 0.001* |

Asterisks indicate that the p-value is smaller than 0.05 after familywise error corrections.

**Figure 4.**
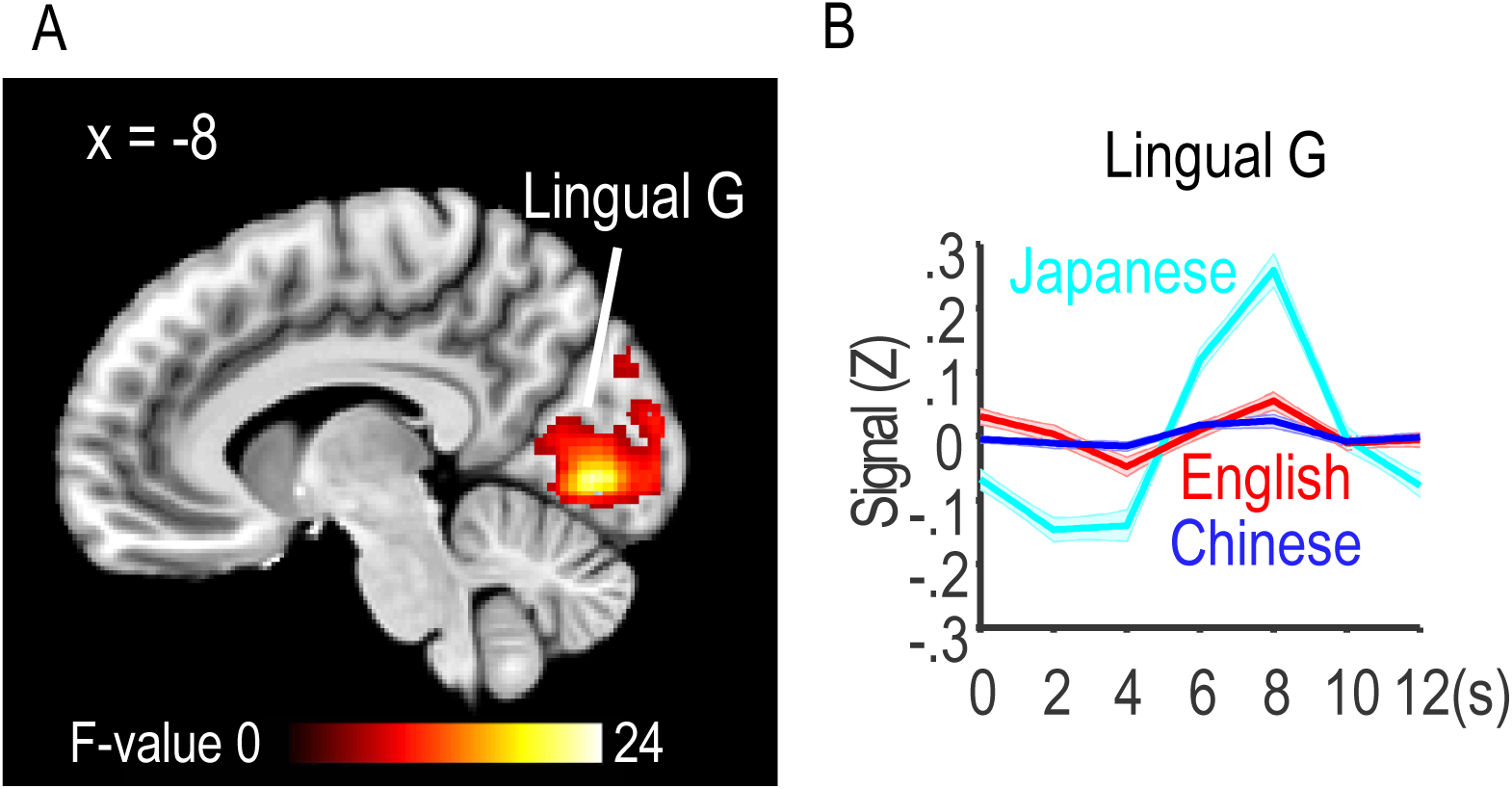
A region that showed a significant effect of language (Japanese, English, Chinese). (A) The cluster distributed around the left lingual gyrus (p < .05, cluster-level FWE). (B) The mean responses of the three language groups.

Finally, we compared activations in response trials with those in no-response trials (Table 4, Fig. 5). The contrast yielded a few clusters that were distributed over the bilateral supplementary motor areas (SMA) and pre-SMA (Fig. 5A), bilateral inferior and middle frontal gyri (Fig. 5B, C) and bilateral primary and higher-order auditory cortices (Figs. 5B, C). Activations became greater in the response trials after 6 s and remained until after 12 s (Fig. 5C-F). It is worth noting that these regions included the left Broca’s area and the left gyrus of Heschl but not the precuneus.

**Table 4.**
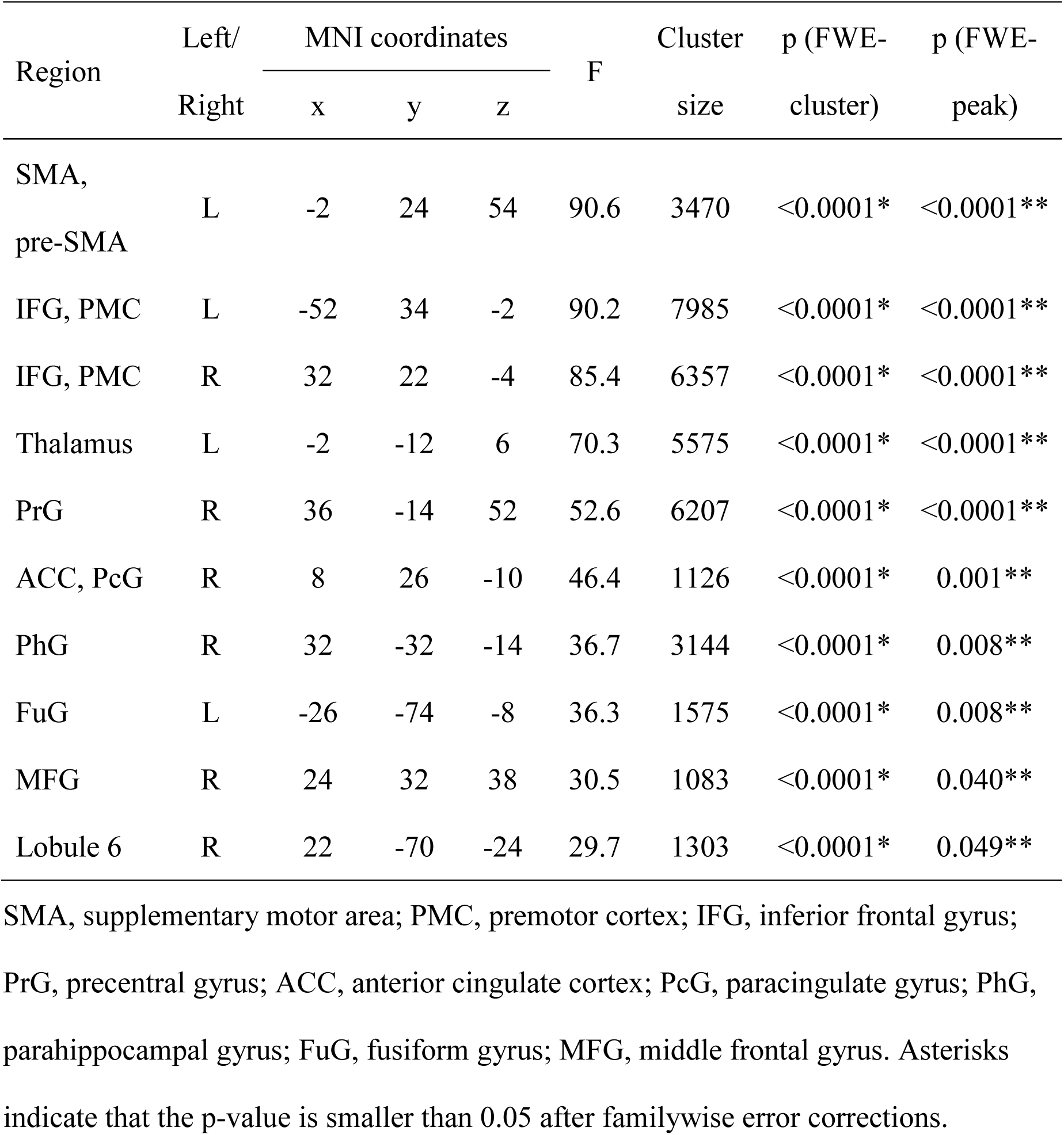
Brain regions that showed a significant main effect of response.

| Region | Left/<br>Right | MNI coordinates |  |  | F | Cluster<br>size | p (FWE-cluster) | p (FWE-peak) |
| --- | --- | --- | --- | --- | --- | --- | --- | --- |
|  |  | x | y | z |  |  |  |  |
| SMA,<br>pre-SMA | L | -2 | 24 | 54 | 90.6 | 3470 | <0.0001* | <0.0001** |
| IFG, PMC | L | -52 | 34 | -2 | 90.2 | 7985 | <0.0001* | <0.0001** |
| IFG, PMC | R | 32 | 22 | -4 | 85.4 | 6357 | <0.0001* | <0.0001** |
| Thalamus | L | -2 | -12 | 6 | 70.3 | 5575 | <0.0001* | <0.0001** |
| PrG | R | 36 | -14 | 52 | 52.6 | 6207 | <0.0001* | <0.0001** |
| ACC, PcG | R | 8 | 26 | -10 | 46.4 | 1126 | <0.0001* | 0.001** |
| PhG | R | 32 | -32 | -14 | 36.7 | 3144 | <0.0001* | 0.008** |
| FuG | L | -26 | -74 | -8 | 36.3 | 1575 | <0.0001* | 0.008** |
| MFG | R | 24 | 32 | 38 | 30.5 | 1083 | <0.0001* | 0.040** |
| Lobule 6 | R | 22 | -70 | -24 | 29.7 | 1303 | <0.0001* | 0.049** |

SMA, supplementary motor area; PMC, premotor cortex; IFG, inferior frontal gyrus; PrG, precentral gyrus; ACC, anterior cingulate cortex; PcG, paracingulate gyrus; PhG, parahippocampal gyrus; FuG, fusiform gyrus; MFG, middle frontal gyrus. Asterisks indicate that the p-value is smaller than 0.05 after familywise error corrections.

**Figure 5.**
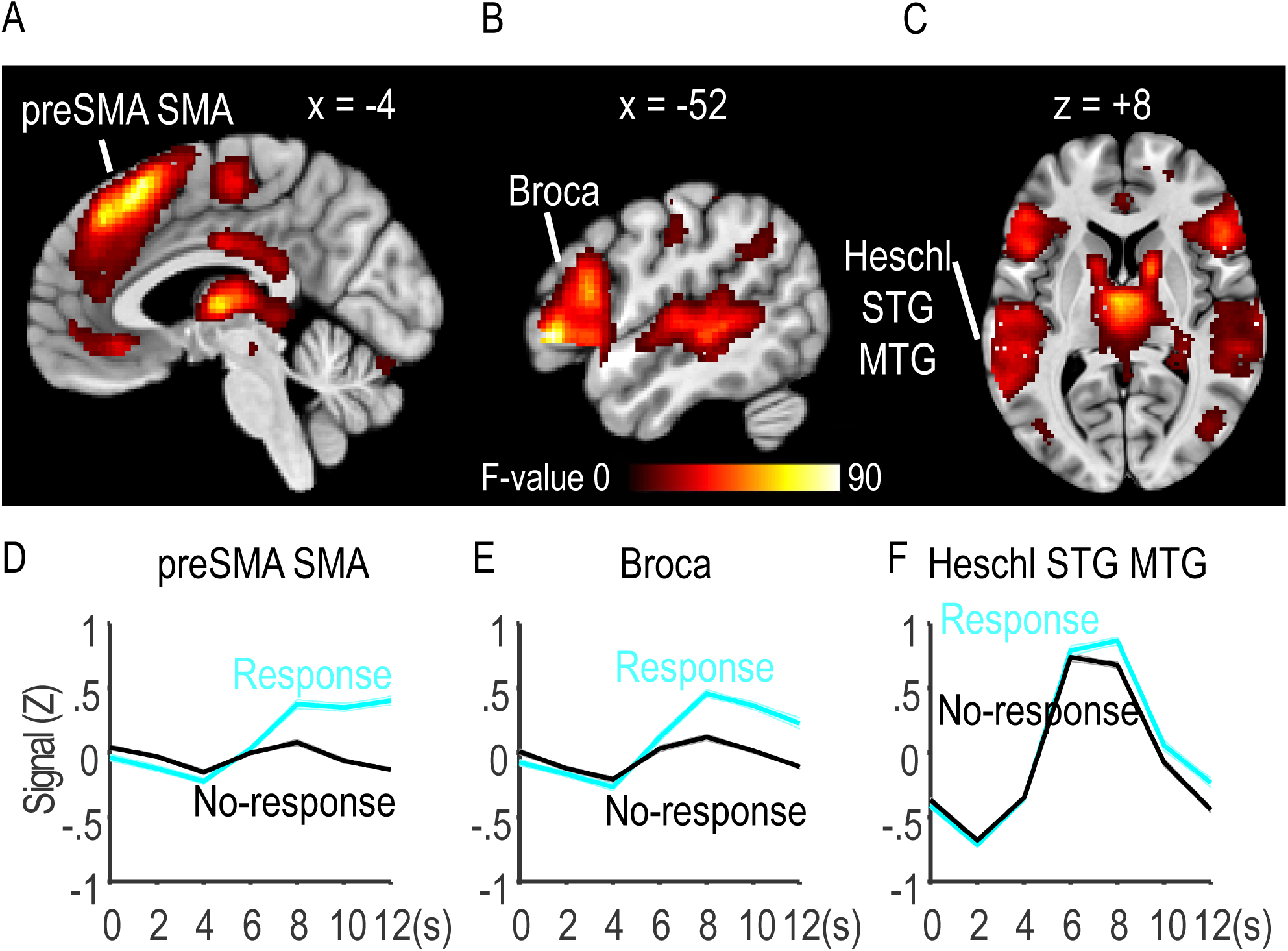
Regions that showed a significant effect of response (response vs no-response). (A-C) A cluster distributed around the supplementary motor area (SMA) and pre-SMA (A). Two other clusters involved Broca’s area (B) and Heschl’s gyrus (C). All clusters were significant after cluster-level FWE correction (p < 0.05). (D-F) The mean responses to unnatural stimuli (response; cyan) and to natural stimuli (no-response, black) in each of the three clusters.

## Discussion

To identify brain regions where the *A* series of time in our mind is represented, we used speech stimuli that would have automatically evoked feelings of “pastness,” “presentness,” and “futurity.” We found three candidate clusters: the precuneus, the left Broca’s area, and the left gyrus of Heschl. However, we should first consider whether all three regions actually represent the *A* series of time in our mind.

First, it may be questioned whether the left transverse gyrus of Heschl could represent the higher concept of time because the region is generally classified as the primary auditory cortex. However, in humans, the left Heschl’s gyrus is larger than the right counterpart. Assuming that this gross anatomical asymmetry represents a morphological substrate of language lateralization (Nieuwenhuys R et al. 2007), the left region in Heschl’s gyrus may have contributed to analyzing phonetic information in intelligible voice stimuli (Scott SK et al. 2000). In agreement with this view, the left Heschl’s gyrus region was activated more strongly and for a longer period by past and future stimuli (Fig. 3F) that were physically longer than the present stimuli (Fig. 2B). Furthermore, the contrast was much enhanced by classifying the stimuli into three groups according to the physical duration of stimuli (short, middle, and long, Table 5, Fig. 6). It is thus probable that the significant difference in activations in Heschl’s gyrus reflected the physical durations of auditory stimuli rather than the perception of time.

**Table 5.** Brain regions that showed a significant main effect of length.

| Region | Left/<br>Right | MNI coordinates |  |  | F | Cluster<br>size | p (FWE-cluster) | p (FWE-peak) |
| --- | --- | --- | --- | --- | --- | --- | --- | --- |
|  |  | x | y | z |  |  |  |  |
| Heschl | R | 50 | -18 | 4 | 204.7 | 3538 | <0.0001* | <0.0001** |
| Heschl | L | -44 | -26 | 10 | 125.5 | 3879 | <0.0001* | <0.0001** |
| PrG | R | 52 | -2 | 50 | 43.2 | 406 | 0.013* | <0.0001** |
| PrG | L | -52 | -4 | 46 | 40.8 | 373 | 0.020* | <0.0001** |
| Insula | L | -40 | 4 | -4 | 19.4 | 394 | 0.015* | 0.004** |
| AnG | R | 34 | -64 | 44 | 15.5 | 1426 | <0.0001* | 0.054 |
| Thalamus | R | 8 | -24 | -2 | 15.5 | 407 | 0.013* | 0.056 |
| SFG | L | -14 | 6 | 70 | 15.0 | 354 | 0.026* | 0.074 |
| IPL | L | -50 | -46 | 48 | 13.2 | 1078 | <0.0001* | 0.241 |
| MCC,<br>PcG | R | 16 | -8 | 42 | 13.2 | 485 | 0.005* | 0.242 |

PrG, precentral gyrus; AnG, angular gyrus; SFG, superior frontal gyrus; IPL, inferior parietal lobule; MCC, median cingulate cortex; PcG, paracingulate gyrus. Asterisks indicate that the p-value is smaller than 0.05 after familywise error corrections.

**Figure 6.**
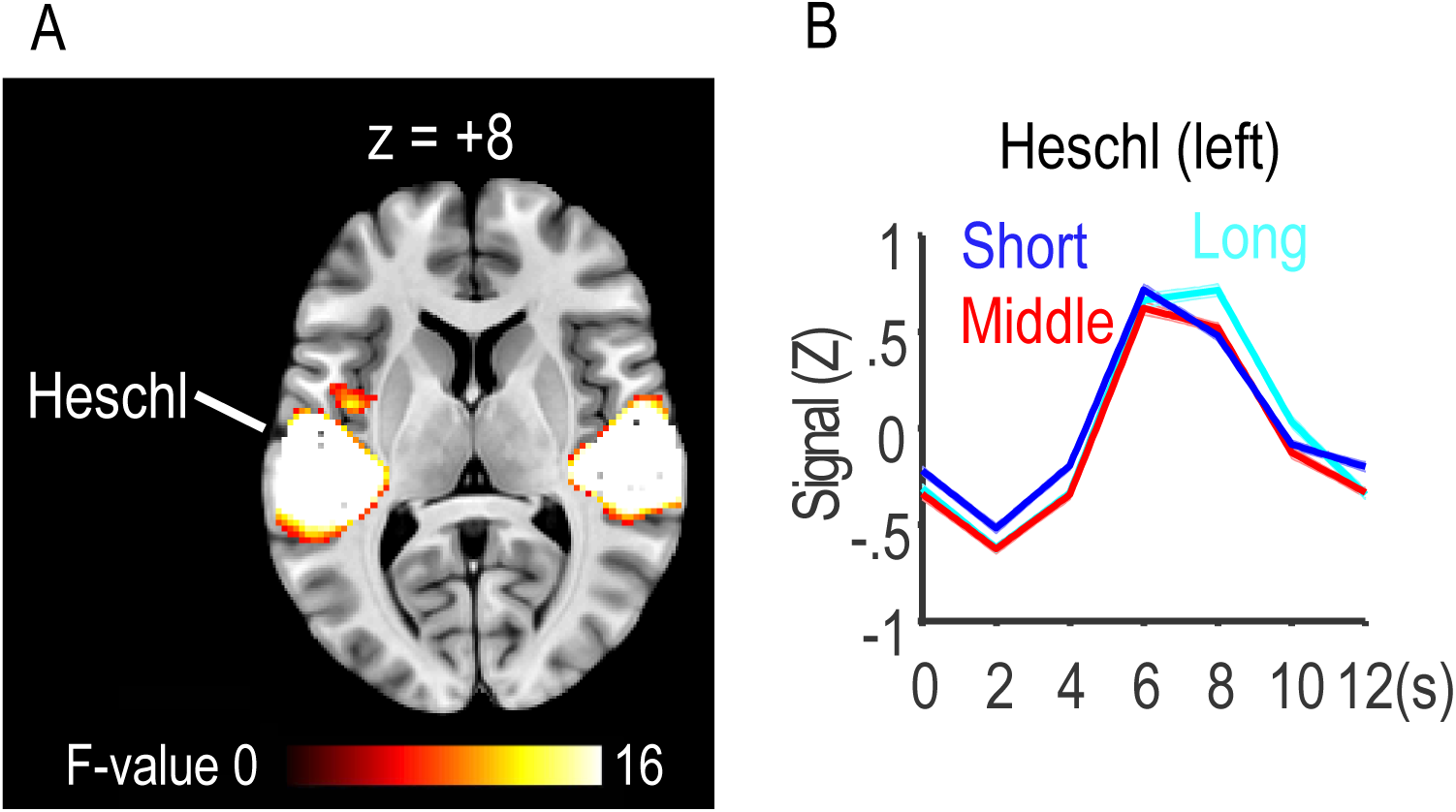
Regions that showed a significant effect of duration of stimuli (short, middle, and long). (A) Two major clusters distributed around the bilateral gyri of Heschl and the temporal plane (p < 0.05, cluster-level FWE). As a result of one-way ANOVA, clusters in Heschl survived after familywise error corrections (p < 0.05). (B) The mean responses (z-scores) in the left cluster. Note that the region was most strongly activated by the long stimuli (cyan) compared to the middle (red) and short (blue) stimuli. Shaded regions show the s.e.m. across 54 participants.

Second, it may be argued that the difference could have been contaminated by the task of judging the naturality of each sentence. We actually found that the left Broca’s area was activated more strongly by unnatural sentences than by natural sentences (Fig. 5). Because the region is known to be activated by silent speech in our mind (Abe K et al. 2011), the difference in activity in Broca’s region could have reflected the difference in the amount of silent speech associated with the judgment of unnaturality. Thus, it was critical to test whether the judgment of naturality in the scanner was independent of the postscan judgment of time. A chi-square test showed that the two factors were independent of each other (chi-square(2) = 2.3, *p* = 0.32). Based on this negative result, we may safely reject the argument that the time-related activations in the candidate regions were contaminated by activations due to judgment of unnaturality.

Finally, it is worth emphasizing that the precuneus was the only region that was free from either argument. It showed 2-3 times greater excitatory responses to the present stimuli than to the past or future stimuli, although the physical duration of the present stimuli was the shortest. Activations in the region did not correlate at all with the judgment of naturality. Thus, these results strongly suggest that the precuneus is involved in representing the *A* series of time and, in particular, the feeling of “presentness”.

### Effects of language

As to the main effect of language, we presently have no idea why the left lingual gyrus was activated more strongly in the Japanese-speaking participants compared with the Chinese- and English-speaking participants. Because the region was shown to be activated during visual imagery even when the eyes were closed (Winlove CIP et al. 2018), the results may suggest that the Japanese participants depended more on visual images in judging whether each sentence was natural or unnatural.

It is worth noting that there were no regions that showed a significant interaction between time and language. The results showed that the precuneus serves as the neural basis of the present in all three languages despite many differences in verb tense systems and word orders.

### Roles of the precuneus in relation to time

It has been repeatedly proposed that a cycle of the alpha rhythm defines a window of simultaneity (Kristofferson AB 1967; Varela FJ et al. 1981; VanRullen R et al. 2005). This function of framing external events in time has not been associated with the precuneus because the source of the alpha rhythm has generally been localized in the posterior visual cortex. However, a recent study (Takahashi T and S Kitazawa 2017) showed that the alpha rhythms consist of five independent components and that the strongest source is located in the precuneus. Thus, it is possible that the precuneus serves as a center that defines a window of simultaneity and the duration of the subjective present.

Our claim that the precuneus represents the present might seem inconsistent with previous findings that the precuneus is important for memory retrieval from the hippocampal formation (Buckner RL et al. 1995; Fletcher PC et al. 1996; Halsband U et al. 1998; Henson RNA et al. 1999; Vincent JL et al. 2006; Raichle ME 2015). These findings have been supported by rich functional and anatomical connections between the precuneus and the hippocampal and parahippocampal networks (Margulies DS et al. 2009). However, we do not think it is a contradiction for two reasons. First, the past stimuli still evoked positive activations in the precuneus (Fig. 3D), although the response was one-third of that to the present stimuli. These previous results showed that clear retrieval of the past evoked stronger responses in the precuneus than when the retrieval was incomplete. Second, memory retrieval is in a sense a re-experience of the past in the present (Addis DR *et al*. 2007; Schacter DL and KP Madore 2016). It is well known that our memory of the past is altered every time we make recall it. A recalled memory could be an experience in the present.

The precuneus is a core region of the default mode network [17]. Interestingly, Raichle ME (2015), a prominent figure in the study of the default mode network, argued against a traditional view that the major role of the precuneus and the default mode network is spontaneous cognition such as mind wandering, daydreaming, and stimulus-independent thoughts (Andrews-Hanna JR et al. 2010). In contrast to the traditional view, he suggested an alternative idea that “the default mode network is playing a critical role in the organization and an expression of preplanned, reflexive behaviors that are critical to our existence in a complex world” (Raichle ME 2015). Our present finding that the precuneus responded most strongly to “presentness”, then to “futurity”, and least to “pastness” could be well explained by this idea of Raichle ME (2015). What is important for our survival and existence is to respond to the clear and present danger. This could be a reason why the precuneus responded most strongly to “presentness”. The precuneus region responded second to the “futurity” because it is worth planning to do something toward an expected event in the future. However, it is of no use to cry over spilt milk. This could be a reason why the precuneus responded least to the past.

Posterior components of the default mode network, including the precuneus and the posterior cingulate cortex, are particularly vulnerable to early deposition of amyloid beta-protein, one of the hallmark pathologies of Alzheimer’s disease (Sperling RA et al. 2009; Dubois B et al. 2014; Palmqvist S et al. 2017). It is interesting that patients with Alzheimer’s disease typically lose orientation in time first, then in place, and then in person (Peer M *et al*. 2015). These clinical observations suggest that the early impairments in the precuneus lead to disorientation in time. This seems to provide evidence for our conclusion that the precuneus region serves as a neural basis of the present, the origin of time, that is required for constructing any orientation in time. The effects of dysfunction in the precuneus on time perception merit further investigation from both basic and clinical viewpoints.

## Acknowledgments

This research was supported by MEXT/JSPS KAKENHI (Grant Numbers 25119001, 25119002, 25119007, 18H05520, 18H05521, and 18H05222) to S.K., T.S. and Y.O. We thank Joseph Tabolt for his assistance in creating the English-language stimuli for this study.

